# Three dimensional parallelized RESOLFT nanoscopy for volumetric live cell imaging

**DOI:** 10.1101/2020.01.09.898510

**Authors:** Andreas Bodén, Francesca Pennacchietti, Ilaria Testa

## Abstract

The volumetric architecture of organelles and molecules inside cells can only be investigated with microscopes featuring sufficiently high resolving power in all three spatial dimensions. Current methods suffer from severe limitations when applied to live cell imaging such as long recording times and/or photobleaching. By introducing a novel optical scheme to switch reversibly switchable fluorescent molecules, we demonstrate volumetric nanoscopy of living cells with resolution below 100 nm in 3D, large field of view and minimal illumination intensities (W-kW/cm^2^).

---

Optical microscopes primarily convey higher spatial information in the lateral directions due to the intrinsic principles of lens-based imaging. The advent of optical nanoscopy has demonstrated the possibility to extract isotropic axial and lateral information beyond the diffraction limit using interferometry^1–3^ and point spread function engineering^4–6^ approaches in both deterministic and single molecule stochastic switching methods. However, the systems capable of conveying truly super resolved 3D information from within the sample today are not suitable for imaging living biological systems, especially over longer times, either because of too long image acquisition times, photobleaching and/or phototoxicity induced by high illumination intensities or chemical preparation to preserve efficient blinking. Other live cell compatible systems only surpass the diffraction limit of a factor of two^7^, which axially is still above 300 nm, or require complex multi-objective setups^8–10^, which severely limits the usability of the system. To tackle this challenge, we designed a novel illumination pattern with intensity modulation along all three dimensions (Figure 1a-d). The pattern exhibits an array of zero-intensity regions co-aligned with the focal plane of detection with sharp and nearly isotropic 3D-confinement (Figure 1b, Supplementary Note 7). The “zeros” are equally spaced and of the same size making it more suitable for coordinate targeted switching schemes such as in STED^11^ and RESOLFT^12^ nanoscopy than previously presented patterns^8,13^. We used the 3D modulated illumination to switch reversibly switchable fluorescent proteins (RSFP)^14^ from a fluorescent (ON) to a non-fluorescent (OFF) state. Since these states are long-lived and require minimal intensities (W-kW/cm^2^) to be completely populated, this enables gentler imaging and to spread the illumination intensity over large areas^15,16^ while maintaining efficient OFF switching. The design of such parallelized optical schemes covering large fields of view of > 40 × 40 μm^2^ is crucial to speed-up the recording time and increase the throughout^17^.

**Figure 1.**
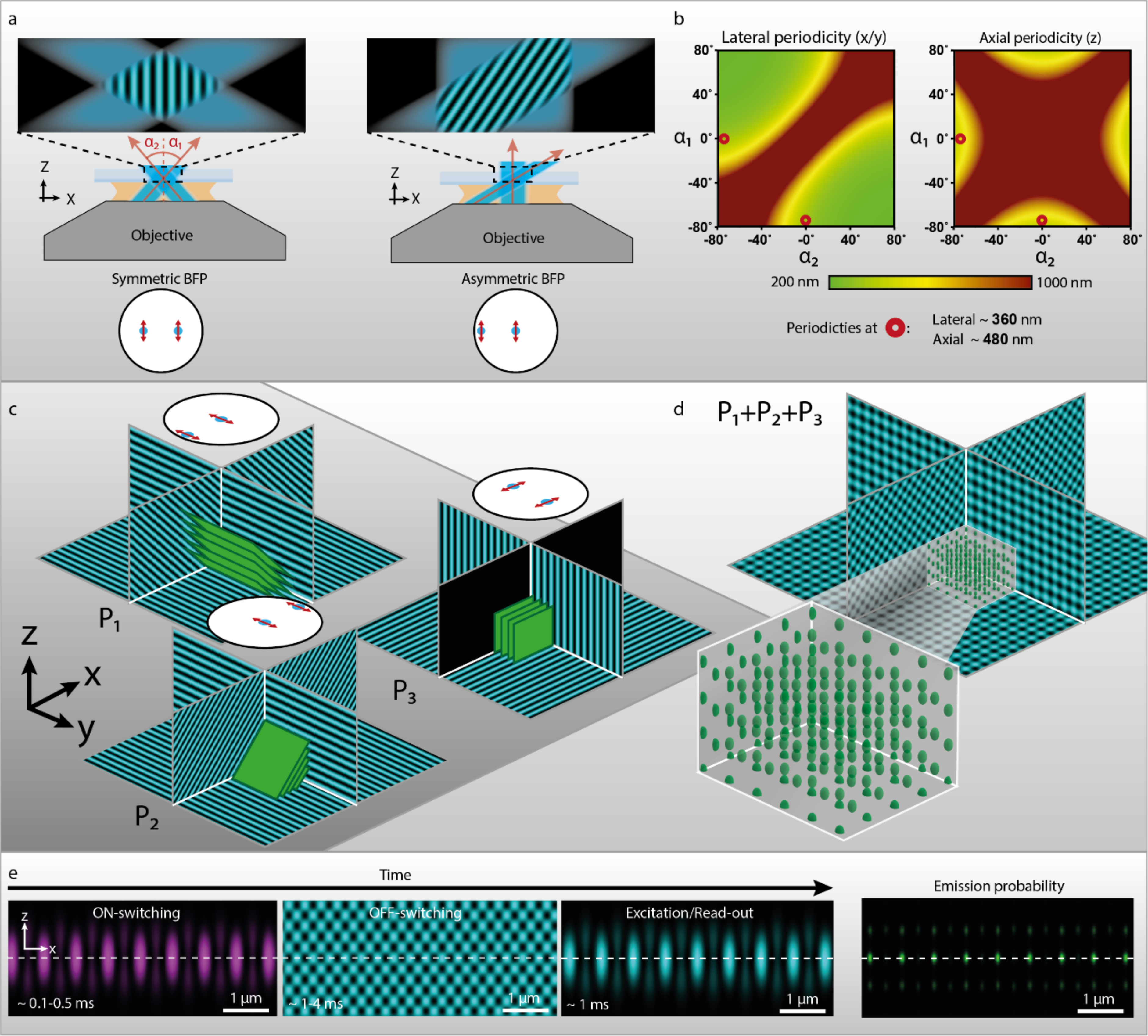
**a)** Schematic illustration of coherent beam interference with different tilt. Below the objectives the corresponding placement of the focused beams on the back focal plane is shown together with their polarization (red double arrows). The tilt of the final pattern depends on the angles α_1_ and α_2_ defined here as the angle between the direction of propagation of the beam and the optical axis (z-axis). **b)**: The 2D-plots show the lateral and axial periodicity of the resulting interference pattern as a function of the angles α_1_ and α_2_. Marked in red circles is the combination of angles used for the tilted patterns in the implemented version of the 3D-pRESOLFT system. **c)** The three interference patterns P_1_, P_2_ and P_3_ are shown with corresponding beam placements on the back focal plane. Green planes illustrate the geometry of zero-intensity planes within each pattern. **d)** The incoherent superposition of P_1_, P_2_ and P_3_ results in three-dimensionally confined volumes emphasized here with green volumes. **e)** Schematic illustration of the imaging sequence used in 3D-pRESOLFT imaging showing simulated illumination patterns in the X-Z-plane in the correct temporal order and duration. The rightmost image shows the calculated expected emission distribution during read-out resulting from the full sequence of illuminations.

The theoretically unlimited 3D resolution of our system is fully owed to the new OFF-switching pattern which is created using the incoherent superposition of three different standing wave patterns. Each standing wave pattern creates planes of zero intensity in the illuminated volume as shown in Figure 1c. The points where planes from all three patterns intersect will be the centers of the resulting zero intensity volumes (Figure 1d). Each standing wave pattern is generated by the interference of two coherent plane waves, exiting the objective at predefined directions. Two of the standing wave patterns, P_1_ and P_2_, are tilted with respect to the optical axis. This tilt gives the final pattern its axial modulation. The tilt is achieved by shifting the two focused spots on the back focal plane so that they are asymmetrically positioned with respect to the optical axis (Figure 1a). The third pattern, P_3_, which is symmetric on the back focal plane, confines the volumes in the final lateral dimension. The exact axial and lateral periodicities of the pattern can be tuned by changing the position of the foci on the back focal plane. Since the periodicity of the pattern in a given direction is inversely proportional to the fluorescence confinement at a certain intensity, tuning the periodicities will affect the properties of the imaging system. For the imaging demonstrated here, we chose a configuration that minimized the axial periodicity and thus maximizes the potential axial resolution. In our configuration, the axial and lateral periodicities are 480 nm and 360 nm respectively.

When combined with RSFPs such as rsEGFP2^14^, the blue-light induced OFF-switching enables us to imprint patterns of state distributions into the sample which then, under fluorescent excitation, translates into a spatial pattern of expected emission (Figure 1e). In our optical scheme, a multifoci pattern at 405 nm is used for switching on any RSFPs located at the focal spots. The sharp zero regions of the OFF-switching pattern are used to sharply confine the ON-state RSFP population and create a pattern of emission consisting primarily of distinctly separated but sharply 3D-confined regions located in the focal plane of the microscope. As these regions are probed with a second multifoci illumination, we can quantify the relative fluorophore density at that coordinate in the sample.

We measured the effective OFF-switching pattern (Figure 2a) by scanning fluorescent beads embedded in Mowiol, where the measured periodicity of 680 nm matches theoretical predictions. Due to the mismatch of the refractive indices between the glass and the sample (1.51 in glass and 1.33 in the sample) the periodicity is compressed from 680 nm in glass/Mowiol to 480 nm in the sample, which gives nearly isotropic confinement. Given the rsEGFP2 switching kinetics and a peak energy of 0.7 J/cm^2^ each for patterns P_1_ and P_2_, the resulting confined fluorescent volume will have a FWHM of around 78 nm (Figure 2b) along **z** and 59 nm along **x**. Adding P_3_ with a peak intensity of 1.4 J/cm^2^ confines the volumes also along **y** to a FWHM of 59 nm.

**Figure 2.**
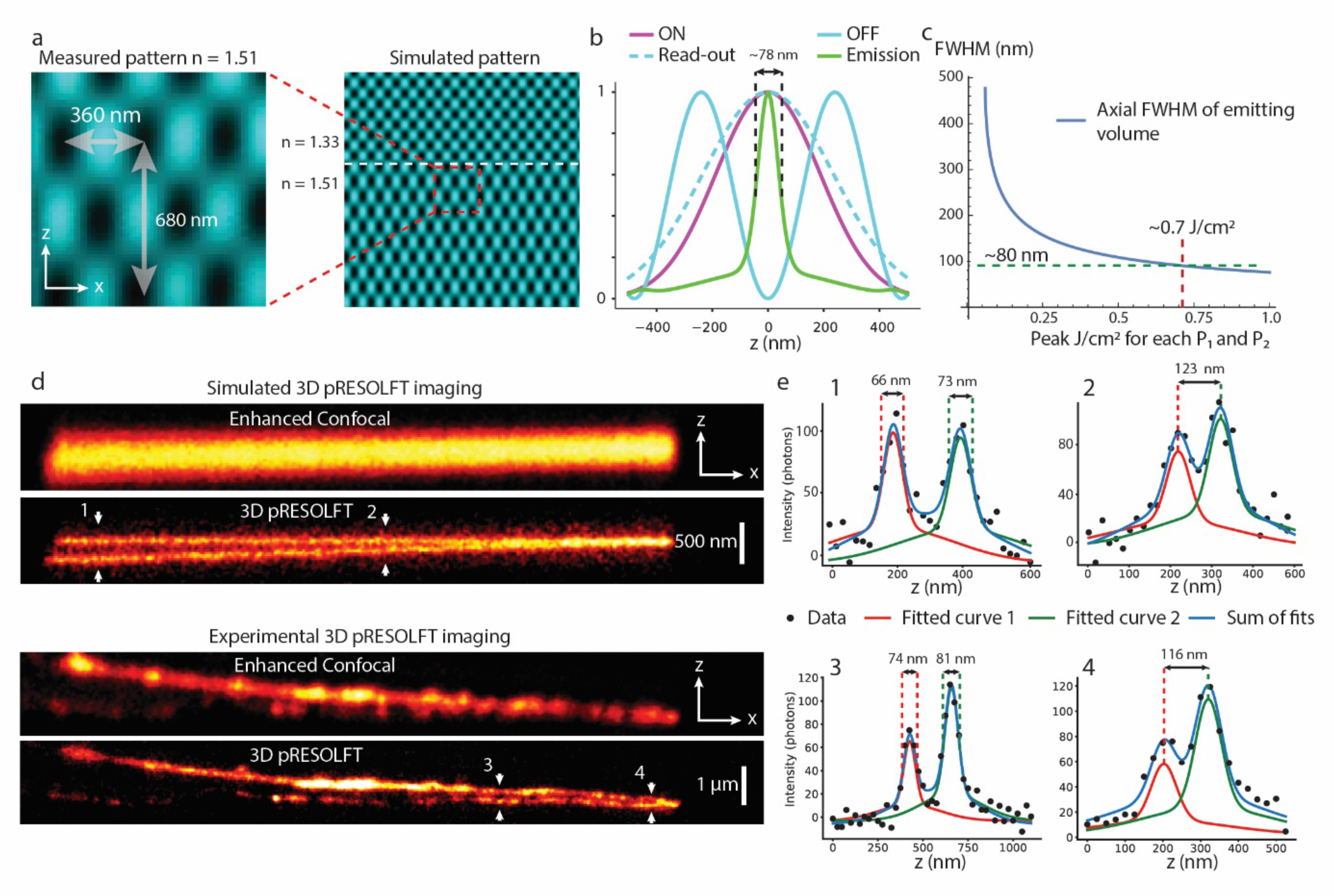
**a)** Left zoom shows the experimentally measured intensity pattern when illuminating the sample with the two tilted interference patterns. On the right the measurement is placed in a larger context showing a simulation of a larger area. Here the effect of a refractive index interface between the cover glass and sample is also illustrated, resulting in an axial compression of the pattern. **b)** The axial line profile of the three different illumination patterns (ON, OFF and read-out) are shown in the left graph as well as the resulting relative expected emission distribution along the same line with the FWHM of the central Gaussian calculated as ~78 nm. **c)** The graph shows the dependence of the central FWHM on OFF-illumination energy where the energy represents the peak energy of pattern P_1_ and P_2_ each. **d)** Panel shows simulated and measured 3D-pRESOLFT imaging. The virtual sample used in the simulations attempt to mimic the structure and labelling of the sample observed in the measured data and is created as two labelled sheets with varying axial separation. The sheets are labelled with 20 fluorophores per 20×20×20 nm voxel. Both measured and simulated data are acquired using only pattern P_1_ and P_2_. To demonstrate confinement with maximal contrast. **e)** Graphs show line profiles along lines labelled in **d**. All line profiles shown are fitted with two the expected effective PSF of the imaging situation, see Supplementary Notes 1 and 2. In the left graphs the widths of the central Gaussians are shows and on the right the separation between the peaks are shown.

The size of the confined volumes describes the potential resolution of the imaging system at those label and imaging parameters. The dependence between axial FWHM and energy of OFF-switching illumination is shown in Figure 2c.

Although the resolution extension beyond the diffraction limit stems from the OFF-switching pattern described above, the final image quality is highly dependent on the multifoci patterns^16^. Adding the multifoci ON-switching and read-out means that fluorophores are only switched on in, and excited from, the confined volumes located in the focal plane of detection. This minimizes the amount of out of focus emission that deteriorates the image SNR, and also gives the flexibility of switching ON and reading out a subset of zero-volumes, which minimizes emission cross-talk and enhances image SNR^16^.

In order to not rely solely on theoretical resolution claims, we focus on practical demonstrations of achievable resolution. To this end we imaged U2OS cells labelled with LifeAct-rsEGFP2 and compare the resulting resolving power with theoretical calculations and computational simulations (Figure 2d-e). The results confirm that the system is able to distinguish axially separated structures at a distance approaching 100 nm. This claim is corroborated by observing a modulation depth of around 60% for structures separated by 116 nm and measuring a FWHM of the confined central Gaussian of around 80 nm (see Supplementary Notes 1 and 2 for details). All values are in good agreement with theory and simulations. As a complementary note, it should be recognized that there is no theoretical band limit to the imaging system, the confined volumes have Gaussian profiles in all directions which intrinsically have infinite frequency support. The amount of high frequency information can be extended by increasing the energy of the OFF-switching light i.e. increasing the confinement. This will however alter the signal to noise and signal to background levels in ways that depend on the fluorophore and labelling characteristics in a non-trivial way. Other parameters like labelling density and type of structure being imaged also alters the effective resolution of the system (Supplementary Figure 6). Deeper analysis of these considerations will help to further increase our understanding of the possibilities and limitations of RSFP based imaging systems.

We further demonstrate the axial resolution improvement by imaging very different types of structures, including notoriously dim and sparse intermediate filament structures such as U2OS cells endogenously expressing Vimentin-rsEGFP2 (Figure 3a-b) as well as the denser and brighter structure of U2OS cells transfected with rsEGFP2-Omp25 labelling the mitochondrial outer membrane (Fig 3c). The dynamics of the mitochondria are also visualized by time lapse imaging of X-Z-slices in Fig 3d, where specifically the axial reorganization can be seen with the high axial resolution of the system. To further explore and demonstrate the imaging ability of 3D-pRESOLFT in living cells we recorded the whole three-dimensional mitochondria network in U2OS cells transfected with rsEGFP2-Omp25 as outer membrane marker. Figure 3e shows the entire volume of two cells recorded with confinement from all three illumination patterns (P_1_-P_3_). In the zooms is highlighted the ability of the system to accurately measure the 3D shape and “hollow” volume of each distinct mitochondria even in closely packed regions such as the nuclear proximity.

**Figure 3.**
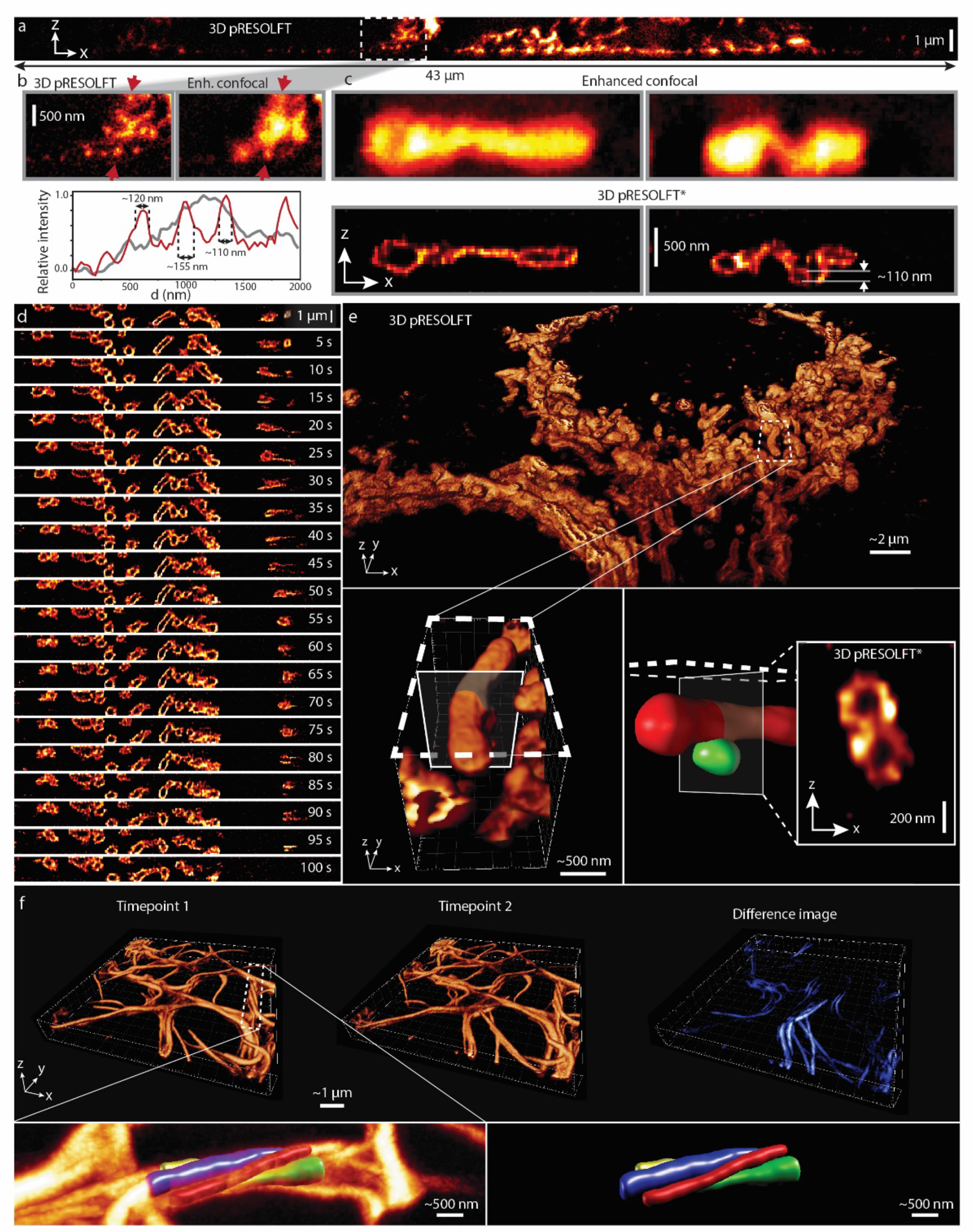
**a)** The full X-Z-slice of a U2OS cell endogenously expressing Vimentin-rsEGFP2 imaged in 3D pRESOLFT mode. **b)** Zoom in of **a** shows a comparison between 3D pRESOLFT and enhanced confocal mode and demonstrates that both the lateral and axial resolution extension is retained also in the notoriously dim endogenously labelled filamentous Vimentin structure. Line profiles plotted in the graph are measured along the line marked with red arrows. 3D pRESOLFT profile shown in red line and enhanced confocal in gray. Both curves are normalized to the maximum intensity of the curve. **c)** Mitochondria labelled with rsEGFP2-Omp25 imaged in both enhanced confocal and 3D pRESOLFT*. The star (*) denotes that the images have been deconvolved as described in the methods section, here using 70 iterations. The images are zooms taken from a bigger slice shown in Supplementary Fig 5. The upper and lower membrane of a small mitochondrial compartment at a distance of ~110 nm are pointed out in the right 3D pRESOLFT* image. **d)** From top to bottom we show an X-Z-slice from a 3D pRESOLFT* (50 iterations) time lapse recording of U2OS cells labelled with rsEGFP2-Omp25 over 2 minutes. The series demonstrated specifically the ability of the system to unveil axial mitochondrial reorganization at high temporal resolution. Images acquired at 5-6 second intervals with frame recording time of 1.7 s. The full recording produces 20 X-Z-slices simultaneously distributed at 720 nm distance in the volume. **e)** Large field of view volumetric image of mitochondrial network in a live U2OS cell labelled with rsEGFP2-Omp25. Zoom in shows a mitochondrial complex in high 3D-resolution. The 3D-resolution enhancement enables the distinction and segmentation of a smaller spherical mitochondrial fragment/vesicle located just beneath the larger one. The volumetric views show raw reconstructed data while the 2D X-Z-slice shown in the bottom right is extracted from a 3D-devonvolved (40 iterations) volume section.

In conclusion, we present a new approach to 3D super resolution imaging with minimal illumination intensities over large fields of view that for the first time in whole living cells enables volumetric studies at resolutions far surpassing the diffraction limit in all three dimensions. The spatial distribution of fluorescent emission from these labels can be finely controlled now in 3D using a novel combination of interference patterns. The technique relies on the use of reversibly switchable fluorophores and is demonstrated here using rsEGFP2^14^ but can be easily applied to other green and red-emitting reversibly switching probes ^18^. The resulting images show both lateral and axial resolutions below 100 nm without prior information or processing. Although robust proof-of-concept demonstrations are provided herein, further development of labeling strategies such as transient binding^19^ has great potential to provide even longer volumetric time lapse imaging. For deeper tissue imaging future implementation of adaptive wavefront corrections^20^ may prove essential for the preservation of the light patterns within these complex media. We believe this technique and future multi-color developments may be of great use for a number of cellular biology studies aiming to dissect the 3D distribution and dynamics of organelles and molecules in living and intact biological systems.

## Acknowledgements

I.T. thanks the ERC starting grant MoNaLISA (http://dx.doi.org/10.13039/501100000781) and the Swedish Foundation for Strategic Research for supporting the project.

## Author contribution

I.T. designed and supervised the project. A.B. engineered and built the microscope with associated software. F.P. prepared the samples. I.T. and A.B. wrote the manuscript with assistance from all the authors.

## Editor’s summary

New optical scheme for parallelized RESOLFT nanoscopy allows volumetric live cell imaging with extended spatial resolution in three dimensions.

## Online methods

### Experimental setup

The 3D pRESOLFT images are acquired using a custom build microscope based on conventional optical and electro-optical components. A schematic of the optical setup is found in Supplementary Figure 7. The optical setup can be divided into three main parts: 1-the two paths to create the multifoci patterns for ON-switching (405 nm laser) and read-out (488 nm laser) 2-The laser paths used to create the OFF-switching pattern (491 nm lasers) 3-The detection path that images the emitted fluorescence onto the sCMOS camera. 1-The two microlenses paths are both generated by a fiber coupled laser source (Cobolt 06-MLD 488 nm and 405 nm filtered with Chroma ET405/10X and ET488/10X respectively) that after a collimating lens passes through the microlens array (Thorlabs MLA150-7AR-M) with a 150 μm distance between lenslets. The image plane after the microlenses is demagnified by a factor of two using a 4f telescope. The beams are then coupled into the main optical path using a 50/50 non-polarizing beam splitter (Thorlabs CCM1-BS013/MB) for the 488 nm path and a dichroic mirror (Semrock Di03-R442-t1-25×36) for the 405 nm path. The main optical path contains an optimized tube lens and objective pair (Leica STED-Orange 1.4 NA Oil objective) giving around 104x demagnification. The final periodicity of the multifoci pattern in the sample is 720 nm.

2) The OFF-switching paths consist of three separate paths originating from two different but identical laser sources (Cobolt Calypso 491 nm DPSS). Using a 50/50 polarizing beam splitter (Thorlabs CCM1-PBS251/M) one of the beams is divided into two to give the total of three beams. Two of them, beam 1 and beam 2, have horizontal polarization and the last one, beam 3, has vertical polarization. The two beams with horizontal polarization are directed onto a diffraction grating (phase-diffraction gratings of 437-nm-high SiO2 lines with a 25 μm period from Laser Laboratorium Göttingen) with horizontal grating lines. These beams will create the partial pattern P_1_ and P_2_ in Figure 1c. The mirrors preceding the diffraction grid are placed so that these beams hit the grid at an angle of 1.125 degrees in opposite directions, Supplementary Figure 7. This small tilt at the plane of the diffraction grids is what gives the asymmetry in the back focal plane. Since these two beams originate from different laser sources, they will be temporally incoherent and thus will not interfere with each other. The beam with vertical polarization is directed onto a diffraction grating with vertical grating lines. This beam is aligned symmetrically with the optical axis and creates the partial pattern P_3_. After passing through the two gratings, the beams are combined with a second polarizing beam splitter. The combined beams pass through a telescope between which a physical mask is placed to block everything except the +1 and −1 diffraction orders of the gratings. Since the −1 order of beam 1 and +1 order of beam 2 will both fall on the optical axis at the plane of the mask and need to be let pass, while the 0 order of beam three should be blocked, a very small intentional misalignment of beam 3 is introduced to allow blocking of the 0 order without blocking the −1 and +1 order of beam 1 and 2 respectively. This will introduce a minor tilt in also the partial pattern P3. This will however have no significant influence on the final pattern and performance of the system. For calculations of final pattern periodicities, see Supplementary Note 7

3) The detection is designed as a conventional widefield microscope. As the main excitation path is reflected off a long pass dichroic mirror (Semrock Di03-R488-t1-25×36) before entering the objective, the emitted fluorescent light will pass through the dichroic mirror and is imaged onto a sCMOS camera (Hamamatsu ORCA-Fusion) using a standard 200 mm tube lens. The detection path also contains an additional bandpass filter (Chroma ET535/70m) to minimize any ambient light and two notch filters (Chroma ZET405NF and ZET488NF) to eliminate any reflections.

### Image reconstruction and processing

In order to reconstruct a final image (2D or 3D) from the raw 3D pRESOLFT data, the emission from each confined volume in each frame needs to be quantified. The array of emitting volumes in the sample in each scan cycle is imaged onto the camera sensor, resulting in an array of diffraction limited PSFs on the sensor. From prior knowledge of the illumination patterns the center of each PSF is known and the camera frame is segmented into sub regions, each containing a single detected PSF. Each sub region is then processed individually and the intensity of the central PSF is quantified by a least squares fitting of a centered Gaussian function on top of a constant background in the same way as was done in our previous work^16^. The final image corresponds to the coefficients associated with the central diffraction limited Gaussian from the fits. The reconstruction algorithm is implemented in Python with GPU-accelerated (CUDA) processing and reconstructs full volumetric data within seconds.

The optical crafting of emission combined with the reconstruction described above results in an image formation model where the final raw reconstructed volumes can be described as a convolution of the underlying sample density with a derived three-dimensional effective point spread function (PSF) with added noise. The effective PSF can be well approximated as a sum of a wider diffraction limited Gaussian function and a narrow super resolved Gaussian, see Supplementary Note 1.4. This type of image formation model is well suited for conventional Richardson-Lucy deconvolution. By accurately calculating the effective PSF for given imaging parameters, the deconvolution allows for significant increases in image contrast at the highest spatial frequencies. The deconvolution is done using a custom script implemented in Python that allows for easy adaptation of the effective PSF to imaging parameters used.

### Sample preparation

U2OS (ATCC® HTB-96™) cells were cultured in Dulbecco’s modified Eagle medium (DMEM) (Thermo Fisher Scientific, 41966029) supplemented with 10% (vol/ vol) fetal bovine serum (Thermo Fisher Scientific, 10270106), 1% Penicillin–Streptomycin (Sigma Aldrich, P4333) and maintained at 37°C and 5% CO2 in a humidified incubator. For transfection, 2 × 10^5^ cells per well were seeded on coverslips in a six-well plate. After one day cells were transfected using FuGENE (Promega, E2311) according to the manufacturer’s instructions. 24–36 h after transfection cells were washed in phosphate-buffered saline (PBS) solution, placed with phenol-red free Leibovitz’s L-15 Medium (Thermo Fisher Scientific, 21083027) in a chamber and imaged.

