## Supplementary for "Three dimensional parallelized RESOLFT nanoscopy for volumetric live cell imaging"

### Supplimentary notes

#### 1 Imaging with reversibly switchable fluorescent proteins (RSFPs)

Reversibly switchable fluorescent proteins switch between two states (ON and OFF) when illuminated with different wavelengths of light. To estimate the probability of fluorophores being in either of the states, we model the system as a two-state Markov system. Different wavelengths of light induce specific switching rates between the two states. As a consequence, patterned (spatially varying) illuminations can be used to create a spatially dependent probability function of fluorophores being in one or the other state. The probability of the two state Markov system being in the ON-state at a certain time can be describe as

$$P(s = ON; t) = \frac{r_{ON}}{r_{ON} + r_{OFF}} + \left(p - \frac{r_{ON}}{r_{ON} + r_{OFF}}\right) e^{-(r_{ON}+r_{OFF})t} \quad (1)$$

Where $r_{ON}$ is the switching rate from OFF to ON, $r_{OFF}$ is the switching rate from ON to OFF, $p$ is the probability of the system starting in the ON-state and $t$ is time.

##### 1.1 Fluorophore properties

The most commonly used fluorophores in RESOLFT imaging are so-called negative switchers, meaning that the absorption peaks for fluorescence excitation and OFF-switching coincide. The fluorophore used herein, rsEGFP2, falls into this category. Using experimental measurements, the different switching rates of rsEGFP2 induced by the specific wavelengths used can be estimated. We see from these measurements that although 488 nm light induces a large OFF-switching rate, the OFF-switching is never fully complete, indicating that under 488 nm illumination, a small residual ON switching rate is still present.

##### 1.2 Switching rate estimation

In order to estimate the different rates induced by the different illuminations, we use a biological sample labelled with the fluorophore of interest, in the scope of this work we limit ourselves to the fluorescent protein rsEGFP2. The sample is first illuminated with 405 nm light to push most of the proteins to the ON-state. Small consecutive pulses of 488 nm light are then applied to the sample and synchronized with the camera in detection. Since only fluorophores in the ON-state are able to emit fluorescence, the amount of fluorescence detected will be proportional to the relative amount of fluorophores in the ON-state. The time trace of detected emission light thus represents the exponential decay of ON-state population, see Supplementary Figure 1. By measuring/calculating the illumination intensity at the sample, we can estimate the decay constant for the given intensity by fitting a function of the form of equation 1 to the decaying fluorescent signal. And by changing the illumination intensity, we can see that the OFF-switching rate seems to scale linearly with increased intensity. Similarly, the rate of ON-switching is measured by pulsing the 405 nm laser and probing the relative emission from a trailing 488 nm pulse. For this measurement we use a 4.3 W/cm2 488 nm pulse that only switches about 3% of the ON-state fluorophores back off with each pulse. When extracting ON-rate, we disregard this minor OFF-switching effect. Under 405 nm illumination, we measure a 72% activation after 1 ms illumination with 252 W/$cm^2$ intensity, Supplementary Figure 2. From the results above we conclude that the ratios between switching rates and intensity are fairly steady. With a good estimate of the switching rates we can calculate how the probability of fluorophores residing in one or the other states evolves over time under a spatially varying illumination intensity.

##### 1.3 Propagation of probabilities

Using equation 1 we can calculate the probabilities of different states after certain illumination sequences. Inserting the illumination patterns used for ON and OFF-switching we can calculate the spatially dependent probabilities $P_{ON}(x, y, z)$, describing the probability of a fluorophore residing in the ON-state if it is located at coordinate $(x, y, z)$. Since we use negative switchers, the fluorophores will undergo switching also during the read-out phase of the illumination sequence. The total expected emission of a fluorophore must therefore be calculated by first calculating the total expected time that the fluorophore resides in the ON-state during the read-out illumination, and then multiplying with the rate of fluorescence emission resulting from the read out illumination intensity, giving

$$T_{ON}(t) = \frac{r_{ON}t}{r_{ON} + r_{OFF}} + \left(p - \frac{r_{ON}}{r_{ON} + r_{OFF}}\right)\frac{1 - e^{-(r_{ON}+r_{OFF})t}}{r_{ON} + r_{OFF}} \tag{2}$$

$$Em(t) = r_{fl}T_{ON} \tag{3}$$

Where $T_{ON}$ is the expected total time in the ON-state, $Em(t)$ is the expected number of emitted photons and $r_{fl}$ is the rate of fluorescent emission under given illumination. Using these equations we can calculate the expected emission from different spatial coordinates that are exposed to different illumination intensities. In Supplementary Figure 3 we show the relative light intensities and probability distributions along an axial line passing through the intensity zero of the OFF-switching pattern. In these calculations we use illumination intensities that attempt to mimic the illumination applied in a typical high resolution 3D pRESOLFT imaging scheme.

Given the form of equation 2 and 3, the expected relative emission can be well approximated as a sum of two Gaussian functions, a wider one resulting from the background emission due to the small residual ON-switching under 488 nm illumination, and a narrow one resulting from the spatial confinement of ON-state fluorophores in the zero-intensity region of the OFF-switching pattern. The accuracy of this approximation is illustrated in Supplementary Figure 3d, where the accurately simulated data is shown to be well approximated by the sum of two Gaussian functions of FWHM values 401 nm and 78 nm respectively.

##### 1.4 Image formation model

We derived above the expected emission function that describes the relative expected number of emitted photons from a given spatial coordinate. Due to the selective activation and read-out the spatial separation between emitting regions is well above the optical diffraction limit of the system allowing for independent quantification of each emitting volume. The quantification is done by segmenting the camera frame into sub-regions centered on the emitting volumes. Digital pinholing is then done by pixelwise multiplication between a pinhole function and the acquired image and the resulting value is assigned to the corresponding volume element (voxel) in the raw reconstructed volume. The transformation between the sample density and the raw reconstructed volume is thus described as

$$V(\mathbf{r}) = \iiint_{\mathbf{u} \epsilon R^3} D(\mathbf{r}) h(\mathbf{u} - \mathbf{r}) d\mathbf{u} + \varepsilon \qquad \mathbf{r}, \mathbf{u} \ \epsilon \ R^3 \tag{4}$$

Where $D(\mathbf{r})$ is the fluorophore density at point $\mathbf{r} \ \epsilon \ R^3$ and $h(\mathbf{u})$ is the three-dimensional impulse response (or apparent/effective point spread function) of the imaging system and $\varepsilon$ is an error due to noise. The three-dimensional impulse response of the system is calculated by first convolving each plane of the detection PSF of the system with the two-dimensional pinhole function used in the quantification step and then multiplying with the relative expected emission function given by the illumination scheme.

$$G(\mathbf{r}) = \iint_{\mathbf{v} \epsilon R^2} PSF_{Det}(\mathbf{r}) P(\mathbf{v} - \mathbf{r}) d\mathbf{v} \qquad \mathbf{v} \ \epsilon \ R^2 \tag{5}$$

$$h(\mathbf{r}) = Em(\mathbf{r}) G(\mathbf{r}) \tag{6}$$

Where $PSF_{Det}(\mathbf{r})$ is the detection point spread function, $P(\mathbf{v})$ is the pinhole function and $Em(\mathbf{r})$ is the relative expected emission from point $\mathbf{r}$ with a given pulse scheme and illuminations.

We can approximate the detection PSF as a Gaussian in all planes with z-dependent width

$$PSF_{Det}(\mathbf{r}) = Exp\left(-\frac{r_x^2 + r_y^2}{2\sigma_{PSF}(r_z)^2}\right) \tag{7}$$

The pinhole function is a set to be Gaussian minus a constant [1]

$$P(\mathbf{v}) = Exp\left(-\frac{v_x^2 + v_y^2}{2\sigma_P^2}\right) - A \tag{8}$$

Meaning that $G(\mathbf{r})$ will in each plane be a Gaussian with larger width minus a constant

$$G(\mathbf{v}) = Exp\left(-\frac{r_x^2 + r_y^2}{2\sqrt{\sigma_{PSF}(r_z)^2 + \sigma_P^2}}\right) - B \tag{9}$$

With the approximation of $Em(\mathbf{r})$ as a sum of a wide and a narrow Gaussian, we can conclude that $h(\mathbf{r})$ will resemble $Em(\mathbf{r})$, again being a sum of two Gaussian with the width of each Gaussian scaled according to

$$Exp\left(-\frac{x^2+y^2}{2\sigma_1^2}\right)Exp\left(-\frac{x^2+y^2}{2\sigma_2^2}\right)=Exp\left(-\frac{x^2+y^2}{2\left(\frac{\sigma_1^2\sigma_2^2}{\sigma_1^2+\sigma_2^2}\right)}\right) \tag{10}$$

Thus $h(\mathbf{r})$ is also well approximated by a sum of two Gaussians, where each Gaussian will be slightly narrower than in $Em(\mathbf{r})$. However, evaluated for the emission functions resulting from our illumination scheme, we see that the shrinking effect on the small central Gaussian of $Em(\mathbf{r})$ will be on the order of 1-2% axially and around 3-5% laterally. The wider Gaussian however, will be shrunk by approximately 10-15% both laterally and axially.

### 2 Line fitting of data

Given the image formation model in equation 4, we see that the emission function acts as the effective PSF in our system in the sense that we expect out final 3D image to be the 3D convolution of the underlying label distribution with the emission function plus noise. When performing curve fitting on line profiles, we assume certain properties of the underlying structure and image formation model and fit a set of parameters to the given model. The theoretically expected line profile however, will depend on the three-dimensional structure of the sample. For a point source, the line profile will simply be the line profile through the three-dimensional PSF. For line or sheet structures, the expected line profile will be projections through the PSF. When fitting functions to line profiles, we naturally want to fit with the expected PSF of that image. By simulating the X-Z image of a single very densely labelled slice we can readily estimate the expected line profile of such structures. We see then that the line profile can also be nearly perfectly approximated as a sum of Gaussians, although in this case, the larger confocal Gaussian is stronger compared to the thinner diffraction limited Gaussian. The height of the larger Gaussian is measured to be 38% of the height of the thinner Gaussian. In Figure 2d-e, we model the dual membrane structure seen as two sheets and thus fit the line profiles using the above mentioned effective PSF i.e. a double Gaussian function. For these curve fits, we fix the ratio of the Gaussians to 0.37 and the width of the large Gaussian to 430 nm. We let the width of the small Gaussian vary as well as the center of all double Gaussian functions. It should be noted here that the exact numbers of primarily the width of the wider Gaussian is very dependent on the exact intensity distribution of the read-out illumination pattern. Above numbers are acquired using simulated multifoci patterns based on scalar interference simulations, ignoring polarization effects. Other simulation approaches might give slightly different numerical results, but the general concept remains the same.

### 3 Richardson-Lucy deconvolution

Given the framework presented in section 1.4 and accurate characterization of fluorophore properties, the image formation model can be concisely described as a convolution between the sample fluorophore density $(D(\mathbf{r}))$)and the derived effective PSF as presented in equation 6. This accurate description of the image formation process allows us to estimate the underlying sample density using a Richardson-Lucy algorithm. The precise description of the effective PSF proves essential to achieve accurate deconvolution results. Attempting to deconvolve with a simplified effective PSF, described only by the central confined Gaussian, gives clearly degraded results as shown in Supplementary Figure 5.

#### 4 Frequency support

We show above that the emission probability function of 3D pRESOLFT can be approximated very well as a sum of Gaussian functions. The super resolution ability of the system comes from the sharp central Gaussian function. Since a Gaussian function, unlike an airy function, has theoretically infinite frequency support, there is no absolute cutoff frequency for which above that, the system carries zero information. Information content will however decrease with increasing frequency and the question of interest is whether the frequency distribution of the information content is sufficient to answer the biological question is at hand. Apart from the shape of the emission probability function, which depends on the switching parameters of the fluorophore, this parameter is heavily influenced by the properties of the labeling i.e. labeling density, label brightness etc., but also on the structure of the sample imaged i.e. sparsity, shape etc. In Supplementary Figure 6, we demonstrate the effect of changing labeling parameters on image quality and effective resolution.

#### 5 Simulations

In order to explore and test different illumination schemes we have developed a tool to simulate imaging data of samples labelled with reversibly switchable fluorophores. A voxelised virtual sample is created containing a certain number of fluorophores in each voxel. Illumination patterns are defined on the same voxel grid as the sample and interactions between the illuminations and the fluorophores follow specific spectral responses preassigned to the fluorophores. The binary states of the fluorophores are tracked during the different illuminations applied using randomly generated numbers and fluorescent emission is calculated according to the applied imaging sequence. The fluorescent emission is virtually recorded on a camera after propagation through an ideal diffraction limited optical system and with camera properties mimicking the camera used in the real optical setup. By simulating our data in this way, we incorporate the true stochastic nature of the fluorophore switching giving realistic simulated data. As shown in Figure 2d, our simulations correspond very well with the recorded data. We can also use the simulations to illustrate the impact of different sample and imaging parameters on the quality of the final data. In Supplementary Figure 6a we illustrate the potential increase in image quality achievable by increasing the labeling density or the label brightness. Due to the stochastic nature of photoswitching, an increased labeling density is preferable to an increased label brightness. These results align previous investigations into photoswitching noise[2]. Although the difference between these images are purely in terms of signal to noise ratio, it is clear that there is a strong coupling between signal to noise ratio and useful resolution. We also demonstrate the concept that sample composition also has an impact on the perceived ability of a system to distinguish different structures. In the upper simulation of Figure 6b using low labeling density the two labeled sheets can just be distinguished, while the single filaments are very hard or impossible to even detect. If labeling density is increased as in the lower simulation of Figure 6b, single filaments can just be perceived while sheets are very well separable.

### 6 Practical considerations

To easy reproduction of our setup and results, we below provide further notes on considerations regarding the optical setup and software.

#### 6.1 Pattern creation

With the 1.4 NA Oil objective used in the current implementation of the 3D pRESOLFT system, laser beams focused on the back focal plane of the objective can exit the objective as collimated beams forming angles between 0-68 degrees to the optical axis. If imaging in media matching the refractive index of oil, this range would define the possible angles of beams available for the pattern creation. When imaging live samples however, the refractive index of the immersion media is commonly very close to that of water. This means there will be a second refraction at the glass-water interface giving an additional refraction of the beam and increasing the angle towards the optical axis. This allows us to theoretically reach angles of 90 degrees after the glass-water interface. At this point though, total internal reflection will occur which is why it is important to stay with beam angles that let the beam propagate into the sample. The rightmost graphs in figure 1a show the resulting lateral and axial periodicities resulting from different combinations of beam angles. It can be noted here that there exists combinations of beam angles that give a perfectly symmetric confinement i.e. the same axial as lateral confinement. Reaching this point however means having to use slightly longer periodicities both laterally and axially. For the images presented here, we instead chose to maximize the achievable axial resolution by minimizing this periodicity, resulting in a slightly anisotropic confinement of around 75When looking at the full OFF-pattern before and after the refractive index interface at the glass water interface, the additional refraction of the beams will result in an axial compression of the pattern. Simulations of this phenomenon is shows in figure 1a. The physical measurement of the pattern shown in the same figure is measured in a media with refractive index 1.51, which is why the measured axial periodicity is 680 nm. After the additional axial compression with a factor of 1.4, the resulting periodicity in water will be around 485 nm.

#### 6.2 Axial scan steps

As mentioned above, the refractive index difference at the glass water interface will cause a refraction of the entering beams. When scanning the sample axially, this refractive index interface will also move causing an axial shift of the OFF-switching pattern in the water. Since the imaging plane is defined by the relative position of the intensity zeros of the OFF-switching pattern with respect to the sample. The step size of the axial scan needs to be set correctly to achieve an isotropic sampling of the image. The correction factor needed to convert physical axial motion of the stage to effective axial scan step is approximately equal to the axial compression factor mentioned above i.e. 1.4, meaning if the stage/sample is moved 50 nm axially, the relative axial motion of the OFF-pattern with respect to the sample is 35.7 nm. This conversion needs to be considered when setting a suitable axial pixel size.

#### 6.3 Image reconstruction

In order to reconstruct a final image from the raw 3D pRESOLFT data, the emission from each confined volume in each frame needs to be quantified. The array of emitting volumes in the sample in each scan cycle is imaged onto the camera sensor, resulting in an array of diffraction limited PSFs on the sensor. From prior knowledge of the illumination patterns the center of each PSF is known and the camera frame is segmented into sub regions, each containing a single detected PSF. Each sub region is the processed individually and the intensity of the central PSF is quantified by a least squares fitting of a centered Gaussian function on top of a constant background in the same way as was done in our previous work [1]. The reconstruction algorithm is implemented in Python with GPU accelerated (CUDA) processing and reconstructs full volumetric data within seconds. Software is available upon request.

##### 6.4 Pattern alignment

The performance of the 3D pRESOLFT imaging system relies on an accurate co-alignment of the three different illumination patterns. This requires both that the periodicities of the patterns match as well as the translational positioning. Matching the periodicities is mainly a matter of designing the optical paths to achieve magnifications that give matching patterns. However, optical components are never 100% perfect (resulting from e.g. chromatic differences) why some manual craftsmanship may be needed to achieve perfect matching. In our case, we achieve this final minor tuning of periodicities using the telescopes following the microlens arrays. By slightly tuning axial positions of these lenses we can tune the final periodicities to exactly equal double the periodicity of the OFF pattern. For the translational positioning (x-y-z overlap), we first align the OFF-pattern to the detection in terms of rotation and axial positioning by tuning the rotation and axial position of the diffraction gratings (mounted on rotation and z-translation mounts). We then tune the rotation and xy-translation of the two multifoci patterns by physically rotating and moving the microlenses on their rotational and translational mounts. The precise co-alignment of the patterns is inspected and tuned using a conventional bead scan procedure, where a fluorescent bead is scanned through the illumination patterns to probe their intensity at different positions.

#### 7 Geometry of 3D OFF-switching pattern

The patterned illumination used for switching OFF the RSFPs in the sample is created by letting several beam pairs interfere, both coherently and incoherently, with each other.

##### 7.1 Interference pattern from two plane waves

A point in the object space of a microscope is defined by its coordinates $x$, $y$, $z$ in a coordinate system with the orthonormal basis $\hat{\mathbf{x}}$, $\hat{\mathbf{y}}$ and $\hat{\mathbf{z}}$. The vector $\hat{\mathbf{z}}$ points away from the objective and the focal plane of the objective coincides with the plane $z = 0$.

Disregarding polarization, the electric field of a monochromatic plane wave passing through the space can be described as

$$E_n(\mathbf{r}, t) = \Re\{A_n e^{i(-\mathbf{k_n}\cdot\mathbf{r}+\omega_\mathbf{n}\mathbf{t})}\} \tag{11}$$

where $\mathbf{r} = (x_r, y_r, z_r)$ is the position vector, $\mathbf{k_n}$ is the wave vector in radians per meter ($|\mathbf{k_n}| = \frac{2\pi}{\lambda}$), $A_n$ is the amplitude of the wave and $\omega_n$ is the angular frequency. For two coherent and equally polarized plane waves $E_1$ and $E_2$ of a given frequency, we have

$\omega_1 = \omega_2 \equiv \omega$ and thus $|\mathbf{k_1}| = |\mathbf{k_2}|$. Where the two waves coincide, the total electric field, $E_{tot}(\mathbf{r}, t)$ will be given by

$$E_{tot}(\mathbf{r}, t) = E_1(\mathbf{r}, t) + E_2(\mathbf{r}, t) \tag{12}$$

Knowing that the sum of same frequency sinusoids gives a new sinusoidal, we can write

$$E_{tot} = \Re\{A_{tot}(\mathbf{r})e^{i\omega t + \phi(\mathbf{r})}\} \tag{13}$$

for some (in the following results irrelevant) $\phi(\mathbf{r})$) and

$$A_{tot}(\mathbf{r}) = \sqrt{A_1^2 + A_2^2 + 2A_1A_2cos((\mathbf{k_1} - \mathbf{k_2}) \cdot \mathbf{r})} \tag{14}$$

If $A_1 = A_2 \equiv A$, the intensity distribution of the resulting field ends up being

$$I_{tot}(\mathbf{r}) = A_{tot}(\mathbf{r})^2 = 2A^2(1 + cos((\mathbf{k_1} - \mathbf{k_2}) \cdot \mathbf{r})) \tag{15}$$

From this equation we can note that $I_{tot}$ will have its highest modulation along the $\boldsymbol{\kappa} = \mathbf{k_1} - \mathbf{k_2}$ vector and will have constant values on any plane to which $\frac{\boldsymbol{\kappa}}{|\boldsymbol{\kappa}|}$ is the normal vector, specifically interesting are the planes with zero intensity, appearing where $cos(\boldsymbol{\kappa} \cdot \mathbf{r}) = -1$ i.e. where $\boldsymbol{\kappa} \cdot \mathbf{r} = \pi + 2\pi n, n \in \mathbb{Z}$

#### 7.2 Incoherent superposition of patterns

We noted above that the interference pattern created by two plane waves results in zero intensity planes periodically distributed in space i.e. confines the zero-volumes in one dimension. To achieve three dimensional confinement of the zero-volumes, we thus need at least three linearly independent patterns. In our first MoNaLISA implementation[1], two patterns were used to confine the zero-volumes in two lateral but ninety degrees rotated dimensions (e.g. $\hat{\mathbf{x}}$ and $\hat{\mathbf{y}}$ dimensions), both these patterns thus fulfill $\boldsymbol{\kappa} \perp \hat{\mathbf{z}}$. In the 3D pRESOLFT implementation, we instead use two patterns that do not fulfill $\boldsymbol{\kappa} \perp \hat{\mathbf{z}}$ to confine the zero-volumes in one lateral as well as the axial dimension ($\hat{\mathbf{x}}$ and $\hat{\mathbf{z}}$) and a third pattern, fulfilling $\boldsymbol{\kappa} \perp \hat{\mathbf{z}}$, to confine in the last lateral ($\hat{\mathbf{y}}$) dimension. We will hereafter index the patterns as pattern $i$ = 1, 2 and 3, following the order in which they were mentioned above. Disregarding the absolute intensities, each pattern can then be fully described by its $\boldsymbol{\kappa}$ vector, hereafter indexed as $\boldsymbol{\kappa_i}$ for pattern $i$.

#### 7.3 $\hat{\mathbf{x}}$ and $\hat{\mathbf{z}}$ confinement

Since we seek to create an array of zero-volumes that is coplanar with the focal plane of the objective (where $\hat{\mathbf{z}}$=0), it is natural to primarily explore cases where $\boldsymbol{\kappa_i}$ for both pattern 1 and 2 lie in the plane spanned by $\hat{\mathbf{x}}$ and $\hat{\mathbf{z}}$ and where $\boldsymbol{\kappa_1}$ and $\boldsymbol{\kappa_2}$ are each others reflections with respect to $\hat{\mathbf{z}}$ (This gives zeros in the focal plane). Since all $\mathbf{k_n}$ have equal magnitude (same wavelength lasers), each $\boldsymbol{\kappa_i}$ maps onto a unique (disregarding permutations) pair of $\mathbf{k_1}$ and $\mathbf{k_2}$. It follows that the pairs of $\mathbf{k_1}$ and $\mathbf{k_2}$-vectors of patterns 1 and 2 must also be reflections of each other. Given these constraints we can describe the incoherent sum of patterns 1 and 2 by the two angles $\alpha_1$ and $\alpha_2$ formed between the $\mathbf{k_n}$-vectors and $\hat{\mathbf{z}}$.

$$\alpha_n = arccos\left(\frac{\mathbf{k_n} \cdot \hat{\mathbf{z}}}{|\mathbf{k_n}|}\right) \tag{16}$$

Letting us also define $\beta$ as the angle between the $\kappa$-vector of a pattern and the $\hat{\mathbf{z}}$-vector. In order to achieve efficient ON-state confinement it is desirable to have a steep intensity gradient surrounding the zero intensity point. To confine efficiently in along both $\hat{\mathbf{x}}$ and $\hat{\mathbf{z}}$ a steep intensity gradient is needed in both directions. It can be shown that the superposition of the two sinusoidal interference patterns discussed above will result in an elliptical confinement in the $\hat{\mathbf{x}} - \hat{\mathbf{z}}$ plane with the amount of ellipticity defined by the ratio between the periodicities of the sinusoidal patterns along $\hat{\mathbf{x}}$ and $\hat{\mathbf{z}}$ respectively. The $\hat{\mathbf{x}}$ and $\hat{\mathbf{z}}$ periodicities can be expressed as

$$p_x = \frac{2\pi}{|\boldsymbol{\kappa}|\, cos(\beta - \frac{\pi}{2})} \tag{17}$$

$$p_z = \frac{2\pi}{|\boldsymbol{\kappa}|\, cos(\pi - \beta)} \tag{18}$$

Allowing us to define the isotropy of the confined region as

$$\frac{p_x}{p_z} = \frac{cos(\pi - \beta)}{cos(\beta - \frac{\pi}{2})} \tag{19}$$

#### 7.4 Additional $\hat{\mathbf{y}}$ confinement

The superimposed patterns discussed in section 7.3 only confines the emitting regions in two dimensions. In order to achieve fully 3-dimensional confinement it is necessary to also confine the emitting regions along the $\hat{\mathbf{y}}$-direction. This is done by superimposing a third interference pattern, $P_3$, with a $\boldsymbol{\kappa}$-vector fulfilling $\boldsymbol{\kappa} \parallel \hat{\mathbf{y}}$. $P_3$ will then add a sinusoidal modulation along the $\hat{\mathbf{y}}$-direction enabling confinement of the emitting region also in the third dimension.

#### 7.5 On circularity and ellipticity

Given equation 15 we can express the intensity distribution of the incoherent superposition of two equally intense interference patterns as (normalizing the maximum intensity to 1)

$$I(\mathbf{r}) = \frac{1}{2}(1 - cos(\boldsymbol{\kappa_1} \cdot \mathbf{r}))) + \frac{1}{2}(1 - cos(\boldsymbol{\kappa_2} \cdot \mathbf{r}))) \tag{20}$$

Where we have shifted the intensity zero to the origin by changing the sign in front of the cosines to minus. In the vicinity of the origin $(1 - cos(x))$ can be well approximated by a parabolic function

$$(1 - cos(x)) \approx \frac{x^2}{2} \tag{21}$$

Thus in the vicinity of the zero intensity it follows that

$$I(\mathbf{r}) = \frac{1}{2}(1 - cos(\boldsymbol{\kappa_1} \cdot \mathbf{r}))) + \frac{1}{2}(1 - cos(\boldsymbol{\kappa_2} \cdot \mathbf{r}))) \approx \frac{(\boldsymbol{\kappa_1} \cdot \mathbf{r})^2 + (\boldsymbol{\kappa_2} \cdot \mathbf{r})^2}{4} \tag{22}$$

$$I_{sat} = \frac{(\boldsymbol{\kappa_1} \cdot \mathbf{r}_{sat})^2 + (\boldsymbol{\kappa_2} \cdot \mathbf{r}_{sat})^2}{4} = \frac{(cos(\alpha_1)\, |\boldsymbol{\kappa}_1|^2\, |\mathbf{r}|)^2 + (cos(\alpha_2)\, |\boldsymbol{\kappa}_2|^2\, |\mathbf{r}|)^2}{4} \tag{23}$$

Where $\alpha_1$ and $\alpha_2$ are the angles between $\boldsymbol{\kappa}_1$ and $\boldsymbol{r}$ and $\boldsymbol{\kappa}_2$ and $\mathbf{r}$ respectively. By introducing $\gamma$ as

$$\gamma = arccos\left(\frac{\mathbf{r}\cdot\hat{\mathbf{z}}}{|\mathbf{r}|}\right) \tag{24}$$

and $\theta$ as

$$\theta = arccos\left(\frac{\boldsymbol{\kappa_1}\cdot\hat{\mathbf{z}}}{|\mathbf{r}|}\right) \tag{25}$$

and enforcing the symmetry between $\boldsymbol{\kappa_1}$ and $\kappa_{\mathbf{2}}$ giving

$$-\theta = arccos\left(\frac{\boldsymbol{\kappa_2}\cdot\hat{\mathbf{z}}}{|\mathbf{r}|}\right) \tag{26}$$

and

$$|\boldsymbol{\kappa_1}| = |\boldsymbol{\kappa_2}| \equiv \kappa \tag{27}$$

we now get

$$I_{sat} = \frac{\kappa^2\,|\mathbf{r}|^2}{4}(cos^2(\gamma-\theta)+cos^2(\gamma+\theta)) \tag{28}$$

$$|\mathbf{r}| = \frac{2\sqrt{I_{sat}}}{\kappa^2}\frac{1}{\sqrt{cos^2(\gamma-\theta)+cos^2(\gamma+\theta)}} \tag{29}$$

We can thus conclude that the distance $|\mathbf{r}|$ to a predefined saturation intensity $I_{sat}$ will have and elliptical shape in the general case of arbitrary $\gamma$, $\theta$ and $\kappa_n$ values with its half axes along $\hat{\mathbf{z}}$ ($\gamma = 0$) and $\hat{\mathbf{x}}$ ($\gamma = \frac{\pi}{2}$). For the special case where $\theta = \frac{3\pi}{4}$ i.e. the $\boldsymbol{\kappa}$-vectors are orthogonal, $|\mathbf{r}|$ will assume a perfectly circular shape. This situation can be achieved in the lateral ($\hat{\mathbf{x}}$-$\hat{\mathbf{y}}$) plane. Supplementary Figure 8 shows that the elliptical shape derived from evaluation of equation 29 corresponds well with the shape of the zero regions in the saturated simulated pattern. The symmetry of the 2D intensity pattern i.e. $\boldsymbol{\kappa_1}$ and $\boldsymbol{\kappa_2}$ are reflections of each other in $\hat{\mathbf{z}}$, means that the intensity profile along $\hat{\mathbf{x}}$ and $\hat{\mathbf{z}}$ will both be pure sinusoidal functions with equal maximum intensity but different periodicity. Thus along these lines we can readily compare the confinement since it will be proportional to the periodicity of the pure sinusoidal and we can define the isotropy as the ratio between the periodicities along the two axes.

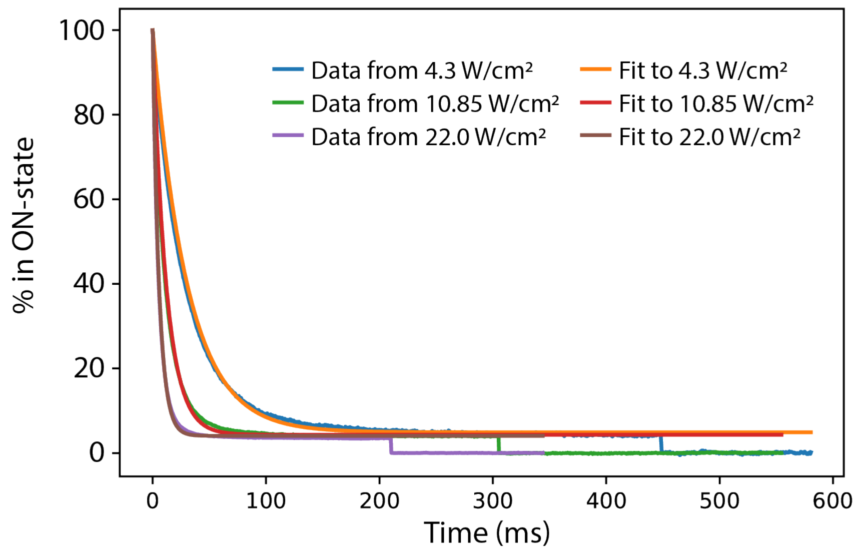


**Supplementary Figure 1:** Graph showing the evolution of ON-state population of rsEGFP2 under three different 491 nm illumination powers and their respective exponential fits. All curves are normalized to the emission detected at the first data point representing the emission from all the proteins starting in the ON-state.

**Supplementary Table 1:** Estimated OFF-switching rates under 491 nm illumination extracted from the fitted curves in Supplementary Figure 1 illustrating the relatively constant ratio between OFF-rate and intensity.

| Intensity of 491 nm illumination [W/$cm^2$] | OFF-rate [events/ms] | OFF-rate/intensity |
|---|---|---|
| 4.3 | 0.0312 | 0.00725 |
| 10.85 | 0.0791 | 0.00729 |
| 22.0 | 0.1685 | 0.00765 |

**Supplementary Table 2:** Estimated ON-switching rates under 491 nm illumination extracted from the fitted curves in Supplementary Figure 1 illustrating the relatively constant ratio between ON-rate and intensity.

| Intensity of 491 nm illumination [W/$cm^2$] | ON-rate [events/ms] | ON-rate/intensity |
|---|---|---|
| 4.3 | 0.00161 | 3.744e-4 |
| 10.85 | 0.00356 | 3.281e-4 |
| 22.0 | 0.00710 | 3.227e-4 |

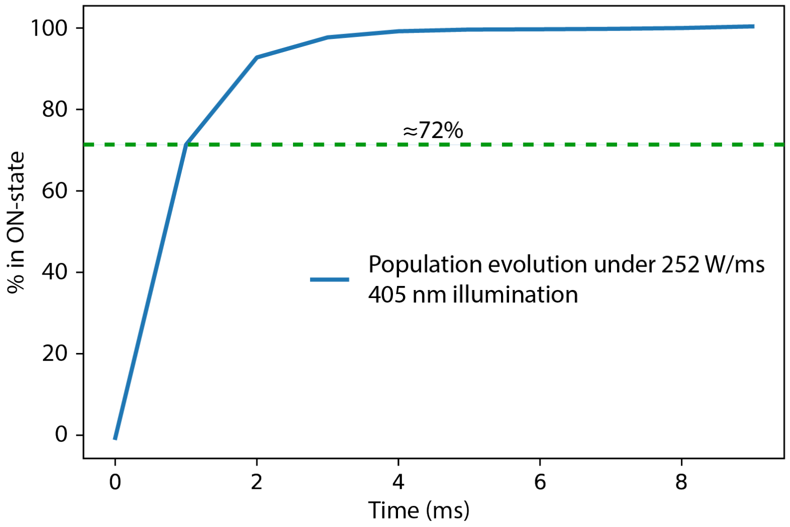


**Supplementary Figure 2:** Graph showing the evolution of ON-state population under 405 nm illumination. Data is acquired by pulsing the 405 nm laser with interleaved 488 nm pulses to probe the fluorescence emission. 488 nm pulses are weak enough to not induce any significant OFF-switching.

**Supplementary Table 3:** Extracted ON-switching rates from the fitted curves in Supplementary Figure 2 illustrating the relatively constant ratio between on rate and intensity.

| Intensity of 405 nm illumination [W/$cm^2$] | ON-rate [events/ms] | ON-rate/intensity |
|---|---|---|
| 252 | 1.3 | 0.0051 |

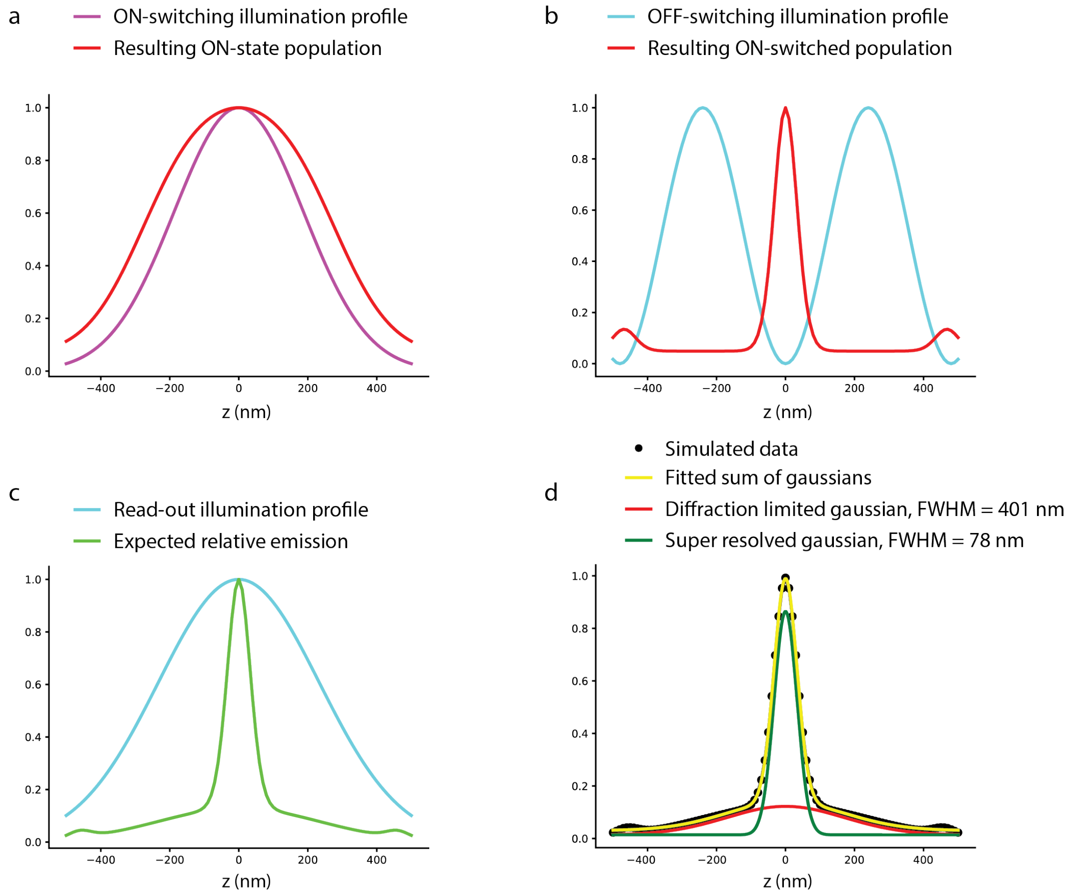


**Supplementary Figure 3:** **a**) Axial illumination profile of ON-switching (405 nm) illumination and resulting ON-state distribution after a 0.192 J/cm2 peak energy pulse. **b**) Axial illumination profile of OFF-switching (491 nm) illumination and resulting ON-state distribution after a 1.424 J/cm2 peak energy pulse. **c**) Axial read-out (488 nm) illumination profile and resulting expected relative emission distribution after 0.188 $J/cm^2$ pulse. All curves in **a-c** are normalized to their individual maximum value. In **d**, simulated data is normalized to its maximum value and fits calculated for the normalized data.

**Supplementary Table 4:** The parameters used to calculate the curves in Supplementary Figure 3 representing a typical 3D pRESOLFT imaging scheme.
*Off-switching rate under 405 nm illumination was not measured. In simulations and calculations of probabilities it is assumed to be zero.

| Illumination [nm] | Peak intensity [$W/cm^2$] | Illumination time [ms] | ON-switching rate [events/ms] | OFF-switching rate [events/ms] |
|---|---|---|---|---|
| 405 | 1920 | 0.1 | 9.8 | 0* |
| 491 | 356 | 4 | 0.12 | 2.55 |
| 488 | 188 | 1 | 0.06 | 1.37 |

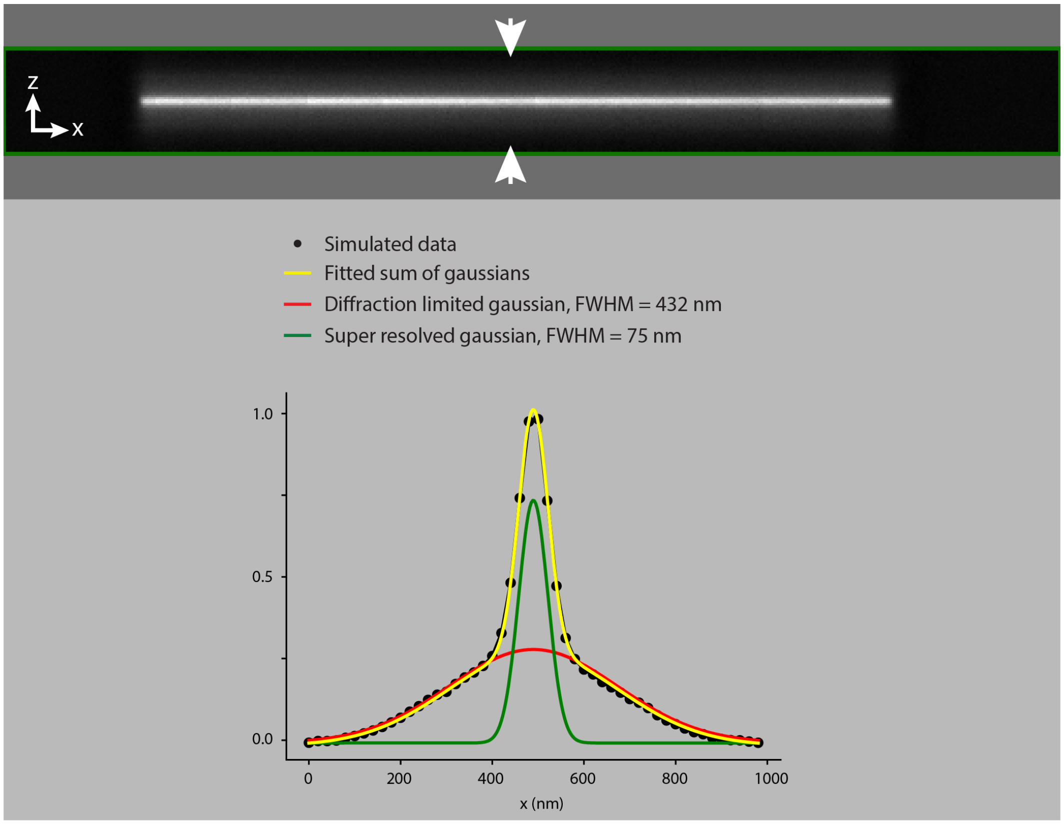


**Supplementary Figure 4:** The expected intensity profile along a line depends both on the 3-dimensional expected emission distribution but also the type of sample being imaged. We here show the resulting axial intensity profile across a virtual sample consisting of a thin sheet, mimicking e.g. a membrane structure.

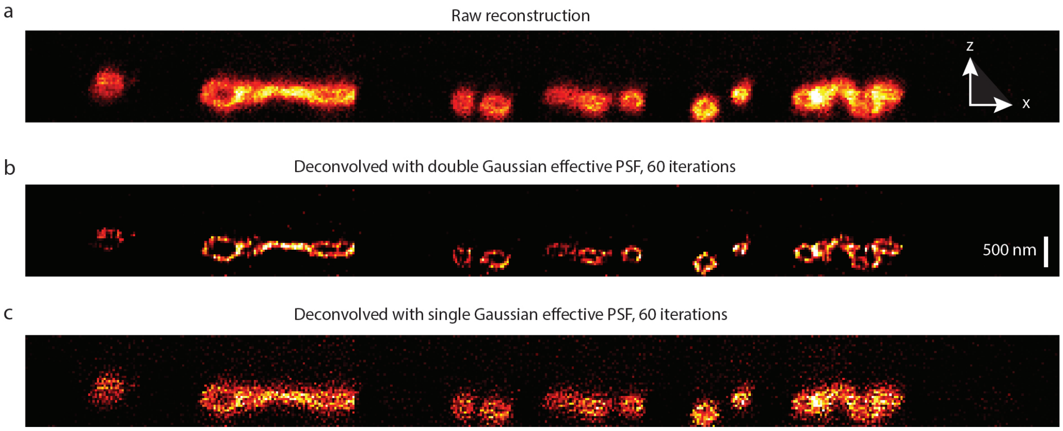


**Supplementary Figure 5:** **a**) The raw reconstructed X-Z-slice from the same sample shown in Figure 3c. **b**) The reconstruction has here been deconvolved using the correct effective PSF as described with our image formation model, consisting of a wider Gaussian plus a narrow Gaussian. **c**) The raw reconstruction has here been deconvolved with the same script as in **b** but with simplified and incorrect effective PSF consisting of only a single sharply confined Gaussian function.

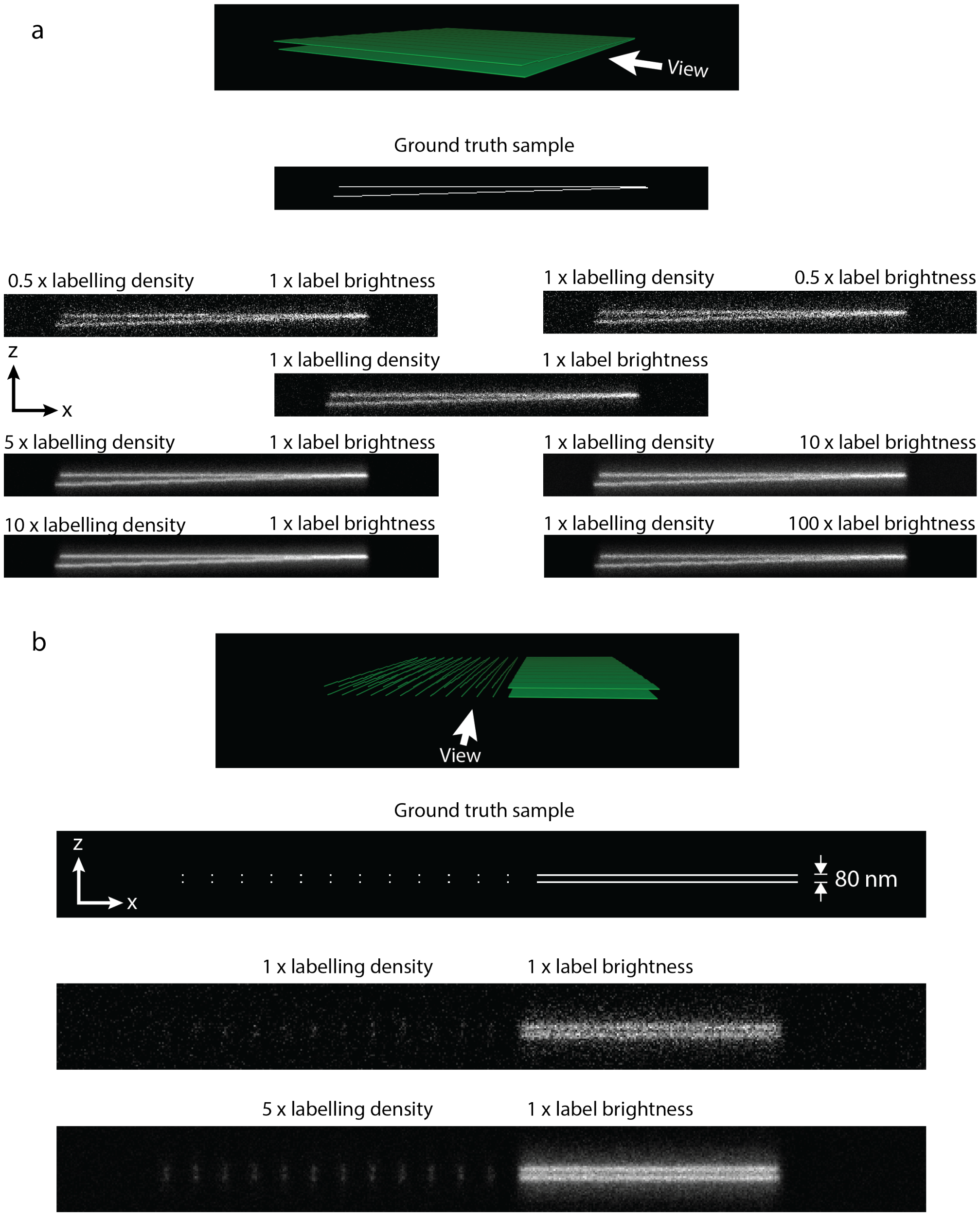


**Supplementary Figure 6:** Demonstration using simulations of how labeling density, label brightness and sample structure affects the practical resolution in RSFP based imaging. Simulations use imaging parameters presented in Supplementary Table 4. **a**) Increasing labeling density does not affect the image quality in the same way as increasing label brightness due to the stochastic nature of photoswitching. Low labeling density with bright labels suppresses the Poisson photon emission noise but does not suppress switching noise. High labeling density however suppresses both noise sources. **b**) At low labeling density, sparse structures like cross-sections of filaments are hardly distinguishable whereas sheet like structures can be both distinguished and separated at 80 nm distance. At 5x higher labeling density, the sparse structure can be clearly distinguished and separated.

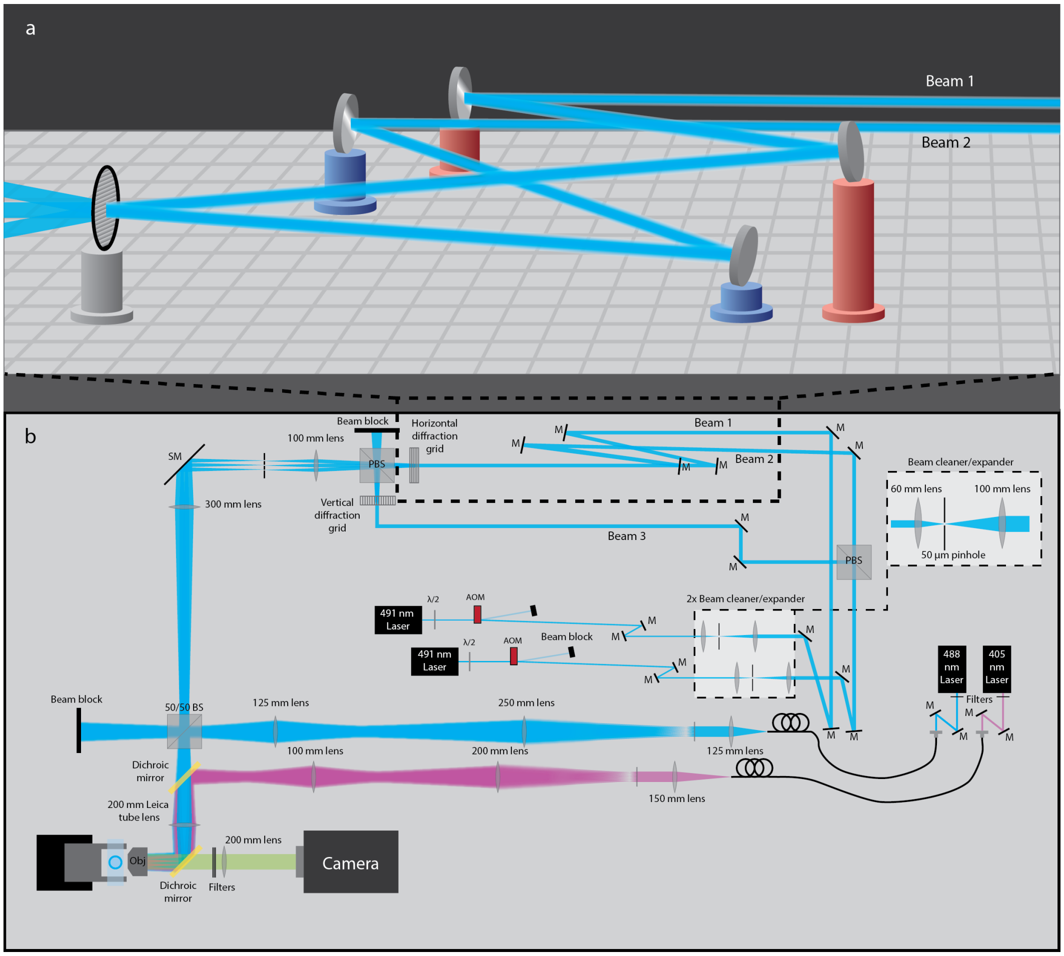


**Supplementary Figure 7:** **a**) Schematic illustration of the optical setup used to acquire 3D pRESOLFT images. BS: non-polarizing Beam splitter, PBS: Polarizing Beam Splitter, M: Dielectric mirror, SM: Silver mirror, Obj: Objective **b**) Three-dimensional illustration of the configuration of optical components used to achieve the asymmetry giving rise to the tilted patterns. Beams are reflected off elevated and lowered mirrors in order to hit the diffractive grating at the correct angle.

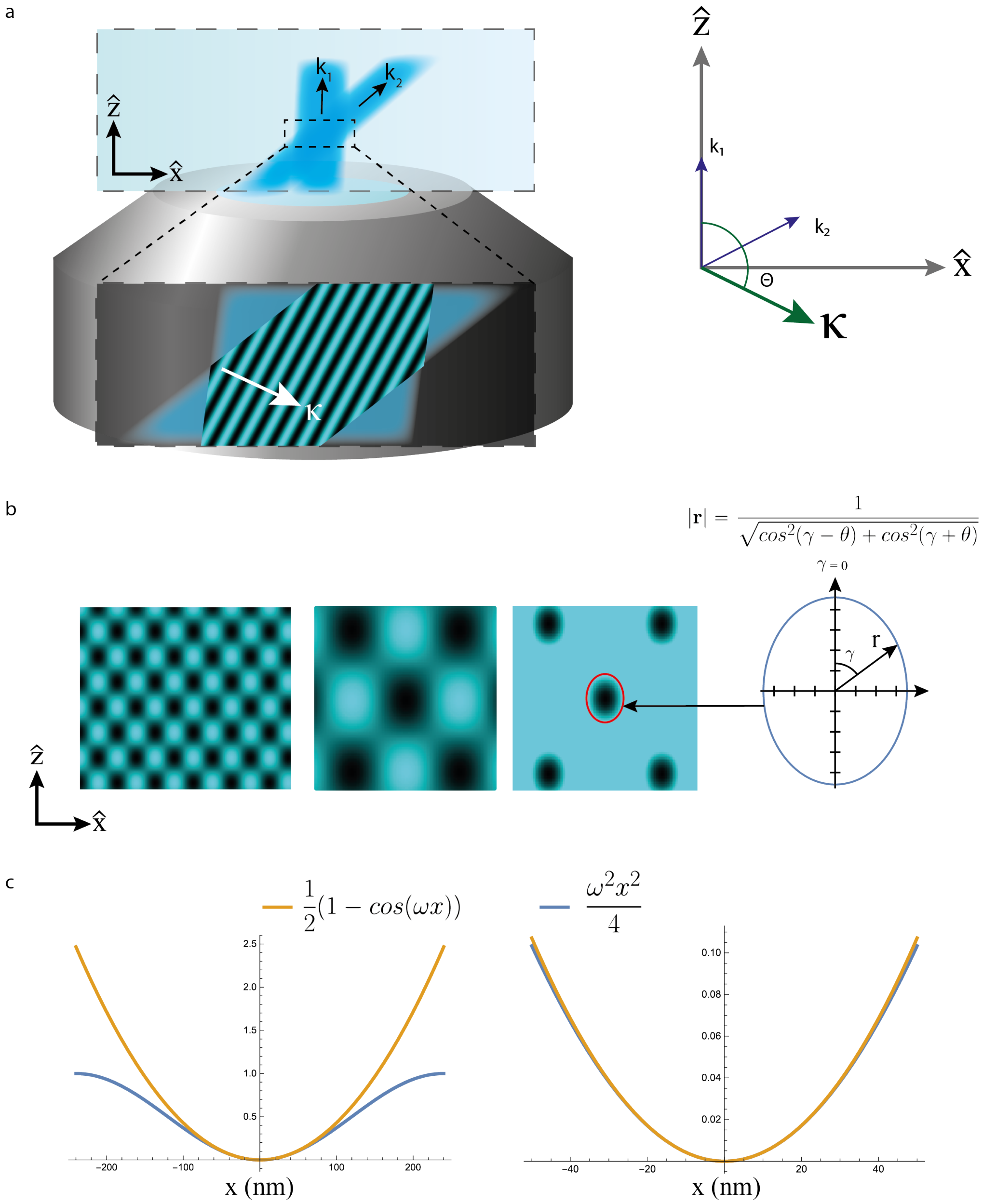


**Supplementary Figure 8:** **a**) Illustration of the geometry described in the section 7.1, showing an example of $\mathbf{k}_n$-vectors and the resulting $\boldsymbol{\kappa}$-vector. **b**) The resulting pattern in the $\hat{\boldsymbol{x}}$-$\hat{\boldsymbol{z}}$-plane from the incoherent superposition of the two tilted patterns $P_1$ and $P_2$. A zoom in on a single zero and a saturated version of the same zoom reveals that the shape of the confined zero matches very well with the shape of the analytically derived ellipse from equation 29. **c**) Graphs showing the wellness of the parabolic approximation in equation 21 for a sinusoidal function in the vicinity of a zero intensity point.

**Supplementary Table 5:** Complete account of imaging parameters used for the experimentally acquired data presented.

| | | ON-switching | | | OFF-switching | | | | | Read-out | | | Scan |
|---|---|---|---|---|---|---|---|---|---|---|---|---|---|
| Fig | Sample | 405 peak intensity (W/cm$^2$) | 405 pulse length (ms) | 405 peak energy (J/cm$_2$) | Confinement dimensions | 491 peak intensity (W/cm$^2$) | 491 pulse length (ms) | 491 peak energy (J/cm$_2$) | X/Y/Z emission confinement (nm FWHM) | 488 peak intensity (W/cm$^2$) | 488 pulse length (ms) | 488 Energy (J/cm$_2$) | Step size (nm) |
| 2 | U2OS Cell labeled with LifeAct-rsEGFP2 | 1920 | 0.5 | 0.96 | X-Z | 356 | 4 | 1.424 | 55/-/78 | 188 | 1 | 0.19 | 25 |
| 3a | U2OS Cell expressing Vimentin-rsEGFP2 | 480 | 0.5 | 0.24 | X-Z | 475 | 2 | 0.95 | 67/-/95 | 251 | 1 | 0.251 | 25 |
| 3c | U2OS Cell labeled with rsEGFP2-Omp25 | 2400 | 0.1 | 0.24 | X-Z | 475 | 2 | 0.95 | 67/-/95 | 188 | 1 | 0.19 | 35 |
| 3d | U2OS Cell labeled with rsEGFP2-Omp25 | 1280 | 0.1 | 0.13 | X-Z | 475 | 1.5 | 0.71 | 76/-/109 | 251 | 1 | 0.251 | 40 |
| 3e | U2OS Cell labeled with rsEGFP2-Omp25 | 960 | 0.1 | 0.096 | X-Y-Z | 949 | 1.5 | 1.423 | 76/76/109 | 251 | 1 | 0.251 | 50 |
| 3d | U2OS Cell labeled with LifeAct-rsEGFP2 | 960 | 0.1 | 0.096 | X-Y-Z | 949 | 2 | 1.90 | 67/67/95 | 251 | 1 | 0.251 | 40 |
| M3 | U2OS Cell labeled with LifeAct-rsEGFP2 | 240 | 0.1 | 0.024 | X-Y-Z | 949 | 1.5 | 1.423 | 76/76/109 | 31 | 1 | 0.031 | 50 |

### References

[1] Luciano A Masullo, Andreas Bodén, Francesca Pennacchietti, Giovanna Coceano, Michael Ratz, and Ilaria Testa. Enhanced photon collection enables four dimensional fluorescence nanoscopy of living systems. doi: 10.1038/s41467-018-05799. URL www.nature.com/naturecommunications.

[2] Benjamin K. Cooper Andrew G. York. Photoswitching noise distorts all fluorescent images. URL https://andrewgyork.github.io/.
